# Genomic heterogeneity inflates the performance of variant pathogenicity predictions

**DOI:** 10.1101/2025.09.05.674459

**Authors:** Baiyu Lu, Xueshen Liu, Po-Yu Lin, Nadav Brandes

## Abstract

Recent studies have reported unprecedented accuracy predicting pathogenic variants across the genome, including in noncoding regions, using large AI models trained on vast genomic data. We present a comprehensive evaluation of these frontier models, showing that performance is inflated by differences in the prevalence of pathogenic variants across genomic contexts. We identify the best-performing models for each variant type and establish a benchmark to guide future progress.

## Background

Mendelian diseases arising from single pathogenic variants account for most of the >7,000 known rare conditions that collectively affect 3-6% of the population^1^. However, current tests provide a clear genetic diagnosis for only ∼50% of cases^2^. This gap largely reflects our limited understanding of variant effects, especially those involving regulatory genomic functions which remain underdiagnosed^3^.

To help close this gap, numerous computational methods have been developed over the past 25 years to predict whether variants are pathogenic or benign. In recent years, a major advance came with large-scale artificial-intelligence (AI) models trained on massive datasets of protein or DNA sequences (**Table 1**)^4-14^. These models not only improved prediction accuracy but, thanks to their largely unsupervised training, also gained greater generality. This approach has proven particularly valuable for regulatory regions, where only a limited number of variants are well characterized and can be used as prediction targets for supervised learning.

**Table 1:** Frontier sequence-based models evaluated in this study.

|  | Sequence modality | Params | Context length | Input | Training data | Training task | Ref |
| --- | --- | --- | --- | --- | --- | --- | --- |
| **Evo2** | DNA | 7B | 8192 | sequence | sequences across species (OpenGenome2) | Autoregressive language modeling | <sup>4</sup> |
| **AlphaGenome** | DNA | 450M | 1M | sequence | functional genomics (e.g., RNA-seq, ChIP-seq) | Predicting 7,058 genomic annotations | <sup>6</sup> |
| **GPN-MSA** | DNA | 86M | 128 | MSA | sequences across species | Masked language modeling | <sup>5</sup> |
| **PhyloGPN** | DNA | 83M | 481 | sequence | sequences across species | Predicting single-nucleotide substitution probabilities from a phylogenetic model | <sup>7</sup> |
| **DNA-BERT2** | DNA | 117M | 512<br>(trained on 128-700) | sequence | sequences across species | Masked language modeling | <sup>9</sup> |
| **Nucleotide Transformer** | DNA | 2.5B | 6000 | sequence | sequences across species | Masked language modeling | <sup>8</sup> |
| **ESM1b** | protein | 650M | 1022 | sequence | sequences across species | Masked language modeling | <sup>13,15</sup> |
| **ESM1v** | protein | 650M | 1022 | sequence | sequences across species | Masked language modeling | <sup>14</sup> |
| **ESM2** | protein | 650M | 1022 | sequence | sequences across species | Masked language modeling | <sup>10</sup> |
| **AlphaMissense** | protein | Unspecified | 256 | sequence, MSA | sequences across species, human allele frequency | Classifying mutations as frequent or absent in humans and primates | <sup>12</sup> |
| **PrimateAI-3D** | protein | Unspecified | Unspecified | 3D protein structure, MSA | 3D protein structure, sequences across species, allele frequency across species | Classifying mutations as frequent or absent in humans and primates | <sup>11</sup> |

MSA: multiple sequence alignment

However, prior studies evaluating these models for pathogenicity prediction have each used different benchmarks and evaluation metrics, making it challenging to directly compare them. Furthermore, existing evaluations have measured performance on highly heterogeneous variant groups. For example, some studies have reported overall accuracy across all noncoding variants, without noting that a substantial portion of labeled variants are at canonical splice sites, which are almost always pathogenic and trivial to detect. As a result, it remains unclear whether these models simply separate broad variant categories (e.g., canonical splice sites vs. other intronic regions) or can also distinguish pathogenic from benign variants of the same type.

To address these gaps, we systematically benchmarked frontier sequence-based AI models (Table 1) at variant pathogenicity prediction and studied their strengths and weaknesses across variant types.

## Results

We curated a benchmark comprising all single-nucleotide variants in ClinVar^16^ labeled as definitively pathogenic or benign, grouped by the types of genomic regions they affect. We benchmarked six DNA-sequence AI models (Evo2^4^, AlphaGenome^6^, GPN-MSA^5^, PhyloGPN^7^, Nucleotide Transformer^8^ and DNABERT2^9^) and five protein-sequence AI models (ESM1b^13,17^, ESM1v^14^, ESM2^10^, AlphaMissense^12^ and PrimateAI-3D^111^) (Table 1). We also included PhyloP^18^, a simple and widely-used phylogenetic model of conservation scores per genomic position, as a baseline (**Figure 1**).

**Figure 1:**
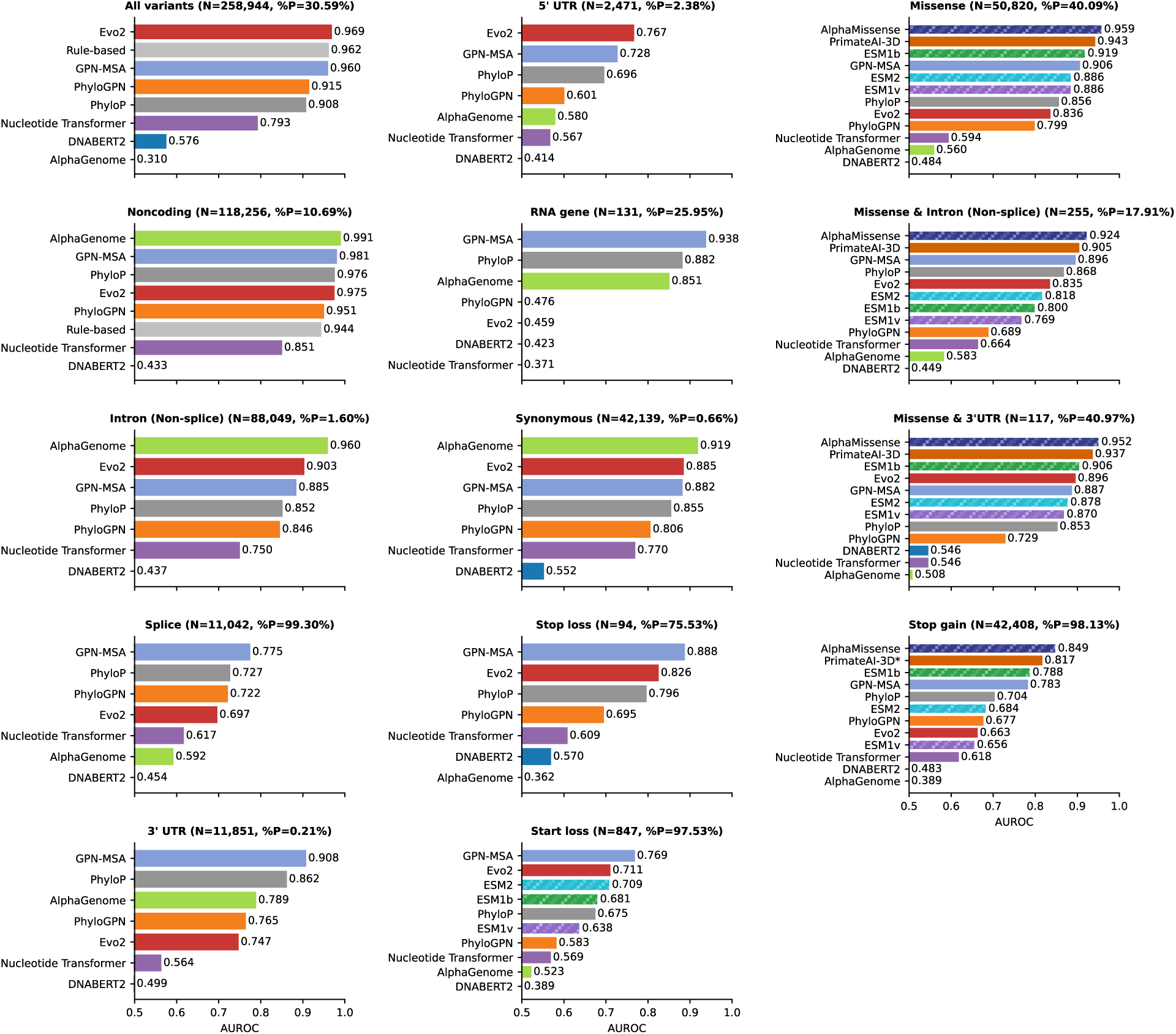
Pathogenicity prediction performance of frontier sequence-based models across variant types. Evaluation and comparison of DNA and protein sequence AI models for their capacity to distinguish between pathogenic and benign variants across variant types, measured by the area under the receiver operating characteristic curve (AUROC). %P indicates the proportion of pathogenic variants in each group. Some groups are defined by multiple annotated effects (e.g., both missense and 3’ UTR, with respect to different transcripts). DNA models are shown as solid bars, protein models as dashed bars. *The evaluation of PrimateAI-3D on stop-gain variants includes only 19,795 variants (see Methods).

All DNA models showed dramatically reduced performance when evaluated within specific variant types. For example, Evo2 appears to perform very well across all noncoding variants (AUROC=0.975), but drops sharply over specific types of noncoding effects (e.g., 0.903 for non-splice intron variants, 0.697 for canonical splice sites). To illustrate the impact of heterogeneous variant groups, we examined the distributions of Evo2 scores across canonical splice and 5’ UTR variants (**Figure 2**). The distribution of Evo2 scores for 5’ UTR variants is shifted towards less damaging prediction for both pathogenic and benign variants, consistent with their much lower prevalence of pathogenicity compared to splice variants (99.3% vs. 2.4%). As a result, merging these two subgroups artificially inflates performance, because pathogenic variants are disproportionately splice variants that receive more damaging scores. Overall, the strong performance of most models observed in the composite groups (all variants and noncoding variants) primarily reflects variant-type heterogeneity. To account for this, we constructed a rule-based baseline assigning each variant a score equal to the frequency of pathogenic variants within its type. This information alone yields strong separation between clinical labels across all variants (AUROC=0.962) and noncoding variants (AUROC=0.944; Figure 1).

**Figure 2:**
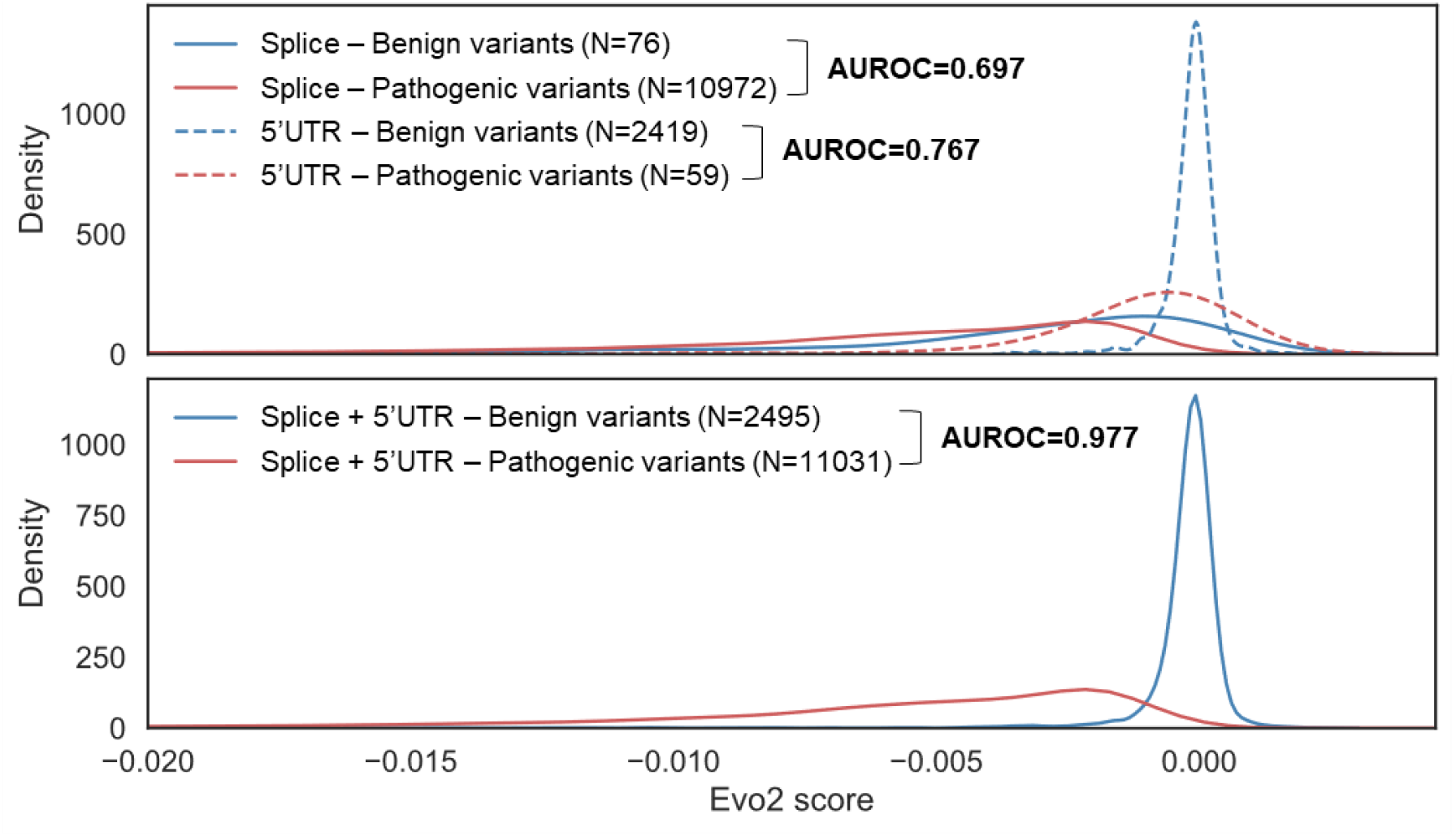
Merging splice and 5’ UTR variants artificially improves the separation between clinical labels achieved by Evo2. Distributions of Evo2 scores across pathogenic and benign variants and across splice and 5’ UTR variants (top) and the union of these two groups (bottom).

We also observe substantial performance variability across different types of noncoding variants, with no single model consistently outperforming the others (Figure 1). For non-splice intronic variants, which comprise 74% of noncoding variants in our benchmark, AlphaGenome performed best (AUROC=0.960), followed by Evo2 (0.903) and GPN-MSA (0.885) that also exceeded PhyloP (0.852). For canonical splice variants, on the other hand, performance dropped across all models, with only GPN-MSA (0.775) surpassing PhyloP (0.727), while AlphaGenome dropped to 0.592. In 3’ UTR variants, only GPN-MSA (0.908) outperformed PhyloP (0.862). For 5’ UTR variants, Evo2 (0.767) and GPN-MSA (0.728) surpassed PhyloP (0.696). For RNA gene variants, GPN-MSA (0.938) again exceeded PhyloP (0.882), but this result is less reliable due to the small sample size (131 variants, 34 pathogenic).

When evaluating performance over coding variants, we also considered protein models, which generally outperformed DNA models (Figure 1). For missense variants, the best performers were AlphaMissense (AUROC=0.959) and PrimateAI-3D (0.943), followed by ESM1b (0.919) and other ESM models. DNA models lagged behind, with GPN-MSA (0.906) coming closest. Interestingly, for variants that are missense in one transcript but noncoding in another (3’ UTR or non-splice intronic regions), protein models showed reduced performance, bringing DNA models closer (but still slightly behind protein models).

For coding variants that are likely to involve DNA- or transcript-level effects beyond amino-acid changes, such as start-loss and stop-gain variants, both DNA and protein models performed worse relative to missense variants. Stop-gain variants were scored by protein models by taking the missense score predicted most deleterious between the mutation site and the end of the protein (see Methods)^17^. This approach worked surprisingly well: AlphaMissense (0.849), PrimateAI-3D (0.817), and ESM1b (0.788) all outperformed GPN-MSA (0.783) and other DNA models. Start loss was treated like missense variants, leading to GPN-MSA (0.769) and Evo2 (0.711) slightly outperforming the ESM models (0.709), but this result is based on only 21 benign variants. Only DNA models were applicable to synonymous and stop-loss variants. For synonymous variants, AlphaGenome performed best (0.919), followed by Evo2 (0.885) and GPN-MSA (0.882) which surpassed PhyloP (0.855). For stop-loss variants (71 pathogenic, 23 benign), GPN-MSA (0.888) and Evo2 (0.826) exceeded PhyloP (0.796).

## Discussion

The metric we used (AUROC) allows direct comparison across variant types despite widely different rates of pathogenicity, ensuring that the observed performance differences can be attributed to model capabilities (see Methods). Our analysis reveals substantial variation in the performance of all models across variant types. No single model was universally optimal, although GPN-MSA and AlphaMissense emerged as the most robust DNA and protein models, respectively, consistently surpassing PhyloP across all variant types. Evo2 and AlphaGenome also led in some contexts, but showed marked drop for other variant types. Notably, AlphaGenome is highly sensitive to the specific method used to extract scores from the model, which may be partly responsible for its highly unstable performance (see Methods). These results suggest a promising meta-prediction approach, where variants are first stratified by type, followed by prediction of the best-performing model within each group.

Our analysis also reveals that combining heterogeneous variant types inflates overall performance, as models can easily infer and exploit variant-type pathogenicity priors, even with limited predictive power within subgroups. These findings underscore the necessity of variant-type-specific benchmarking and choosing models with respect to specific variant types. To help future evaluations meet this standard, we have released our benchmarks and code (see Data and Code Availability).

For certain variant types, the state of the art is highly reliable (AUROC>0.9). This includes missense (AlphaMissense), synonymous (AlphaGenome), non-splice intron (AlphaGenome), 3′ UTR (GPN-MSA) and RNA gene variants (GPN-MSA). By contrast, stop-gain, start-loss, stop-loss, splice and 5′ UTR variants remain difficult for current sequence-based models.

The superior performance of GPN-MSA, AlphaMissense and PrimateAI-3D likely stems from their use of multiple sequence alignment (MSA) inputs, which provide rich evolutionary information. However, reliance on sequence homology makes these models difficult to run locally and limits their generality beyond point mutations. Language models, in contrast, tend to be more general and are also applicable to indels^17^. PhyloGPN takes an interesting middle path, using MSA during training but sequence only during inference. However, it consistently underperforms both the PhyloP baseline and its relative method GPN-MSA. The authors of PhyloGPN note that it relies on a relatively simple evolutionary model (F81)^19^, which may partly explain these results.

This study has several limitations. First, we restricted our analysis to single-nucleotide variants. Second, our evaluation was constrained by the scope of ClinVar, which lacks definitive clinical labels for certain variant types, such as those in promoters or trans-regulatory elements. Third, our analysis of stop-gain variants did not account for nonsense-mediated decay (NMD). Specifically, protein models are expected to perform better on variants not expected to result in NMD (e.g., according to the “50bp rule”)^17^ while DNA models may hold greater potential to capture NMD. However, this likely requires context spanning multiple exons, whereas most DNA models have a relatively short context length (e.g., 8,192bp for Evo2; Table 1).

## Method

### ClinVar benchmark preparation

We downloaded the ClinVar^16^ variant summary file (release: 2025-03-31) and extracted 259,600 single-nucleotide variants mapped to the GRCh38 genome with definitive pathogenic or benign labels (**Supplementary Figure S1**). Each variant was annotated using two complementary strategies. First, variants were mapped to transcripts according to the MANE RefSeq annotation database (v1.4 GFF file).^20,21^ Second, transcript information and amino-acid changes were also extracted directly from the HGVS nomenclature provided by ClinVar, providing an independent annotation source. Variants were categorized as noncoding if neither annotation indicated a protein-coding region. Genomic region categories were assigned from either source when available, including start loss, stop loss, canonical splice site (defined as intron variants within 1-2nt of exon boundaries), and RNA genes. Missense, synonymous and stop-gain categories were taken from the ClinVar HGVS annotations, while the 5′ UTR, 3′ UTRs and non-splice intron categories were derived exclusively from MANE annotations. The final benchmark and code for generating it are available online (see Data and Code Availability).

### Calculating and comparing AUROC scores

We evaluated how well models distinguish between pathogenic and benign variants within each variant type using the area under the receiver operating characteristic curve (AUROC). AUROC has a straightforward probabilistic interpretation: it is the probability that a randomly chosen pathogenic variant receives a more damaging score than a randomly chosen benign variant. For example, an AUROC of 0.9 means that in 90% of such pairs, the pathogenic variant is ranked above the benign one. This makes AUROC an intuitive measure of a model’s ability to separate the two clinical labels. AUROC is also insensitive to class imbalance, since it conditions on the selection of one pathogenic and one benign variant, regardless of their prevalence. This is critical in our study, as variant type categories differ dramatically in pathogenicity prevalence. For example, 99.3% of canonical splice variants are pathogenic compared to 1.6% of all other intron variants. Despite this disparity, the AUROC metric allows direct comparison between these variant types. In addition, AUROC evaluates the full ranking of predictions without requiring calibration of the scores or selection of a cutoff, allowing models to be compared on equal footing. Other metrics, such as accuracy and the area under the precision-recall curve (AUPRC), are sensitive to class balance and therefore unsuitable for this study.

For models in which more negative scores indicate higher pathogenicity (Evo2, GPN-MSA, PhyloGPN and the ESM models), we multiplied the scores by -1 so that higher values consistently indicate greater pathogenicity. We then computed the AUROC using these transformed scores and the pathogenicity labels (pathogenic=1, benign=0). For each variant group (all variants, noncoding variants, or a specific variant type), variants with missing values in any of the compared models were excluded to ensure direct comparison on the same variant set. The only exception was stop-gain mutations, for which only 19,795 out of the 42,408 variants had PrimateAI-3D scores.

### Variant sequence construction

For DNA models, we extracted sequences from the GRCh38 human genome, centered around each variant position, with sequence length determined by the model’s input context (Table 1). For protein models, we used the full amino-acid sequence of the annotated protein isoform. If the protein exceeded the model’s context length, we applied a sliding window approach, subdividing the sequence into overlapping windows of 1,022aa with >511aa overlaps.^17^ The final score was computed as a weighted sum of the scores across the relevant segments.

### Scoring stop-gain variants by protein models

Protein models do not directly provide predictions for stop-gain variants. To estimate their impact, we used the most deleterious prediction of possible missense variants within the truncated protein region (between the introduced stop codon and the end of the protein). For the ESM models, we considered all possible missense mutations, whereas for AlphaMissense and PrimateAI-3D we only considered missense mutations that can result from a single-nucleotide substitution between two codons (as only these scores were available to us).

### Evo2

We used the *evo2_7b_base* model with 8,192bp context length, downloaded from the Evo2 GitHub repository (https://github.com/ArcInstitute/evo2).^22^ Variant scores were computed using the built-in *model*.*score_sequences* function over the reference and alternative sequences separately, to obtain the log likelihood of each sequence. The variant effect score is the log likelihood ratio between these two sequences, calculated as the log likelihood of the alternative sequence minus the log likelihood of the reference sequence.

### GPN-MSA

We used the precomputed GPN-MSA variant scores, downloaded from the authors’ dataset (https://huggingface.co/datasets/songlab/gpn-msa-hg38-scores/resolve/main/scores.tsv.bgz), which provides predicted effect scores for all possible single nucleotide variants in the GRCh38 genome.^5^

### PhyloGPN

We obtained prediction scores from PhyloGPN using the code provided in the model’s official GitHub repository (https://github.com/songlab-cal/gpn/blob/main/examples/phylogpn/basic_example.ipynb). ^7^

### Nucleotide Transformer

We used the Nucleotide Transformer 2.5B Multi-Species model available on HuggingFace (https://huggingface.co/InstaDeepAI/nucleotide-transformer-2.5b-multi-species) with a context length of 6,000 bp.^8^ As suggested by the authors, reference and alternate sequences were processed independently, and the L2 distance between their last hidden layer embeddings at the CLS token was used as the variant effect score.

### DNABERT2

We used the DNABERT2–117M model via HuggingFace (https://huggingface.co/zhihan1996/DNABERT-2-117M).^9^ Because DNABERT2 does not use single-nucleotide tokens, we could not directly compute log likelihood ratios from its logit outputs. Instead, we applied the same method proposed by Nucleotide Transformer. Specifically, we used a context length of 512 bp, which typically corresponds to fewer than 128 tokens under its byte-pair encoding, consistent with the sequence lengths the model was trained on. Reference and alternate sequences were processed independently, and the L2 distance between their last hidden layer embeddings at the CLS token was taken as the variant effect score.

### ESM models

We followed the same pipeline described in our previous publication.^17^ For ESM1b, we used the *esm1b_t33_650M_UR50S* model from the official ESM GitHub repository (https://github.com/facebookresearch/esm).^13^ For ESM1v, we used *esm1v_t33_650M_UR90S_1*, the first of the five models comprising the ESM1v ensemble.^14^ For ESM2, we used *esm2_t33_650M_UR50D*.^10^ All ESM models used a 1,022aa context length. For each variant, the reference sequence was used as input, and the variant effect score was calculated as the difference in logits (log-likelihood scores) between the alternative and reference residues.

### AlphaMissense

We downloaded the precomputed AlphaMissense variant effect scores from the authors’ dataset (https://alphamissense.hegelab.org/), which provides predictions for all possible missense variants in the human genome.^12^ There were 87 variants with more than one AlphaMissense score matched, which we excluded from our analysis of missense variants across all models.

For stop-gain variants, we mapped the RefSeq IDs from our ClinVar dataset to UniProt IDs in the AlphaMissense precomputed score files using the MyGene API.^21,23,24^ We then collected all missense scores within the UniProt sequence at or downstream of the stop-gain position and selected the most deleterious score among them.

### PrimateAI-3D

We downloaded the precomputed PrimateAI-3D variant effect scores from the official model’s website (https://primateai3d.basespace.illumina.com/).^11^ For stop-gain variants, we excluded 22,613 variants whose ClinVar isoform (identified by RefSeq ID) did not match any of the isoforms in the PrimateAI-3D precomputed score file, leaving only 19,795 successfully mapped stop-gain variants used in our evaluation of this model.

### PhyloP

We downloaded the precomputed PhyloP conservation scores based on the 100-way vertebrate alignment under the GRCh38/hg38 assembly from the UCSC Genome Browser.^18,25^

### AlphaGenome

The AlphaGenome model outputs 5,930 human and 1,128 mouse genomic annotations across 11 modalities, including gene expression, splicing, chromatin states, and chromatin contact map. We used the model via the official API (https://deepmind.google.com/science/alphagenome/) with a 1Mbp input context and organism set to human, enabling outputs from all available modalities.^6^ To calculate variant effect scores, we used the recommended implementation, which converts each output to quantiles and assigns variants the maximum percentile observed across all outputs.

Taking the maximum percentile across all outputs, whether or not they are relevant to a given variant type, likely introduced noise to the final scores, as some outputs are more informative than others for predicting pathogenicity. For splice variants, we also considered the splice variant scorer defined in the original publication, which takes a weighted sum of raw scores from the three splice-related modalities (splice site usage, splice site, and splice junction). We also considered scores based on each of the three modalities, maximizing either raw scores or quantiles (**Supplementary Figure S2**). We found that the variant scorer reported in the original publication works better (AUROC=0.708) than the composite score based on all modalities (AUROC=0.614), but that an even simpler scorer based just on the raw scores from the splice site usage modality performs even better (AUROC=0.779). Further work is needed to establish the optimal variant effect prediction strategy for AlphaGenome across different variant types.

## Data and Code Availability

Our full benchmark, which includes the ClinVar labels, genomic annotations and model scores of each variant, is provided as **Supplementary Table S1**. Our source code, which includes the calculation and extraction of predictions from each of the evaluated models, performance calculations (AUROC), and the generation of presented plots, is available as an open-source project at https://github.com/Brandes-Lab/VEP-eval.

---

**Supplementary Figure S1.**
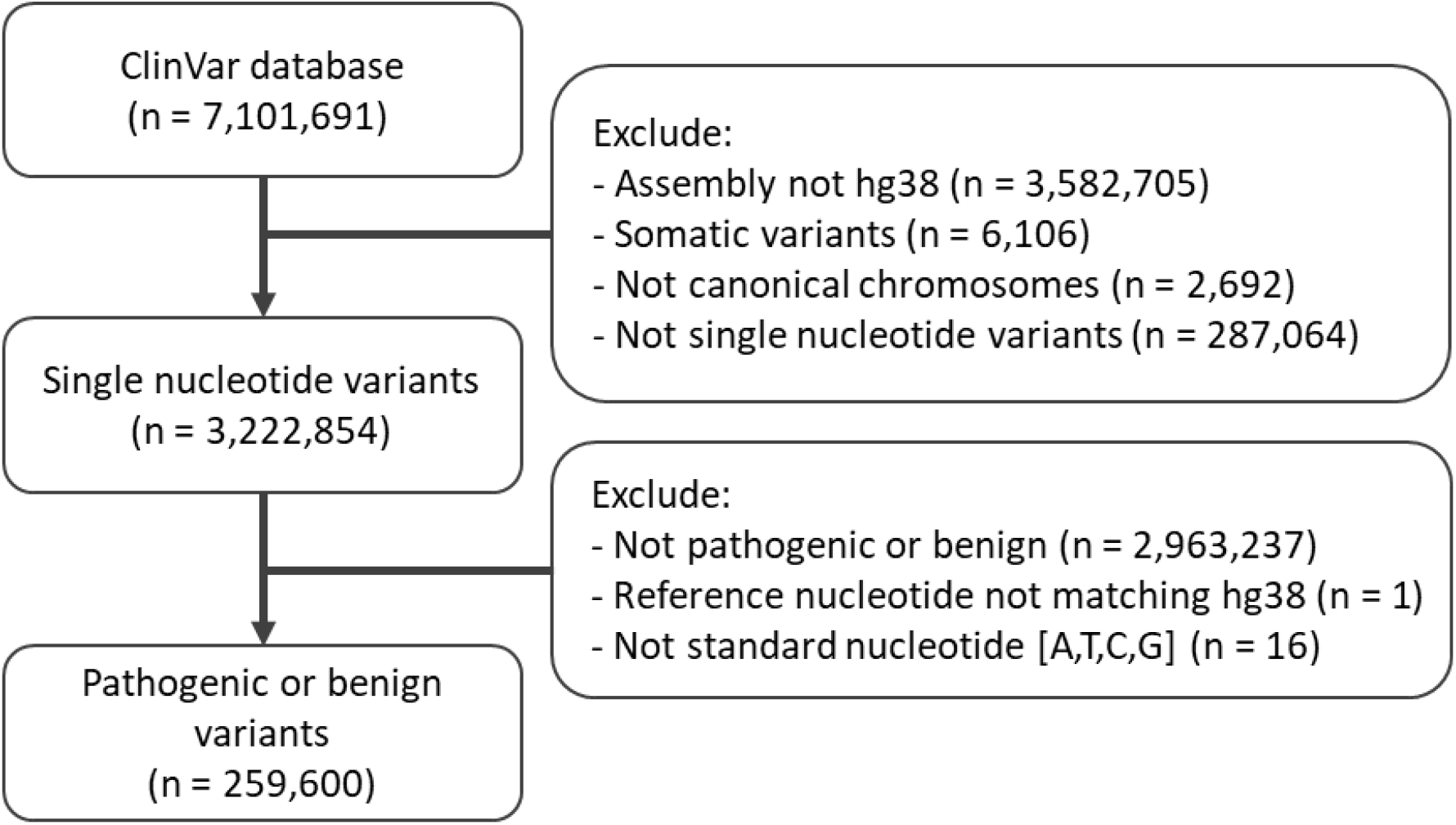
Pipeline for generating the ClinVar benchmark.

**Supplementary Figure S2.**
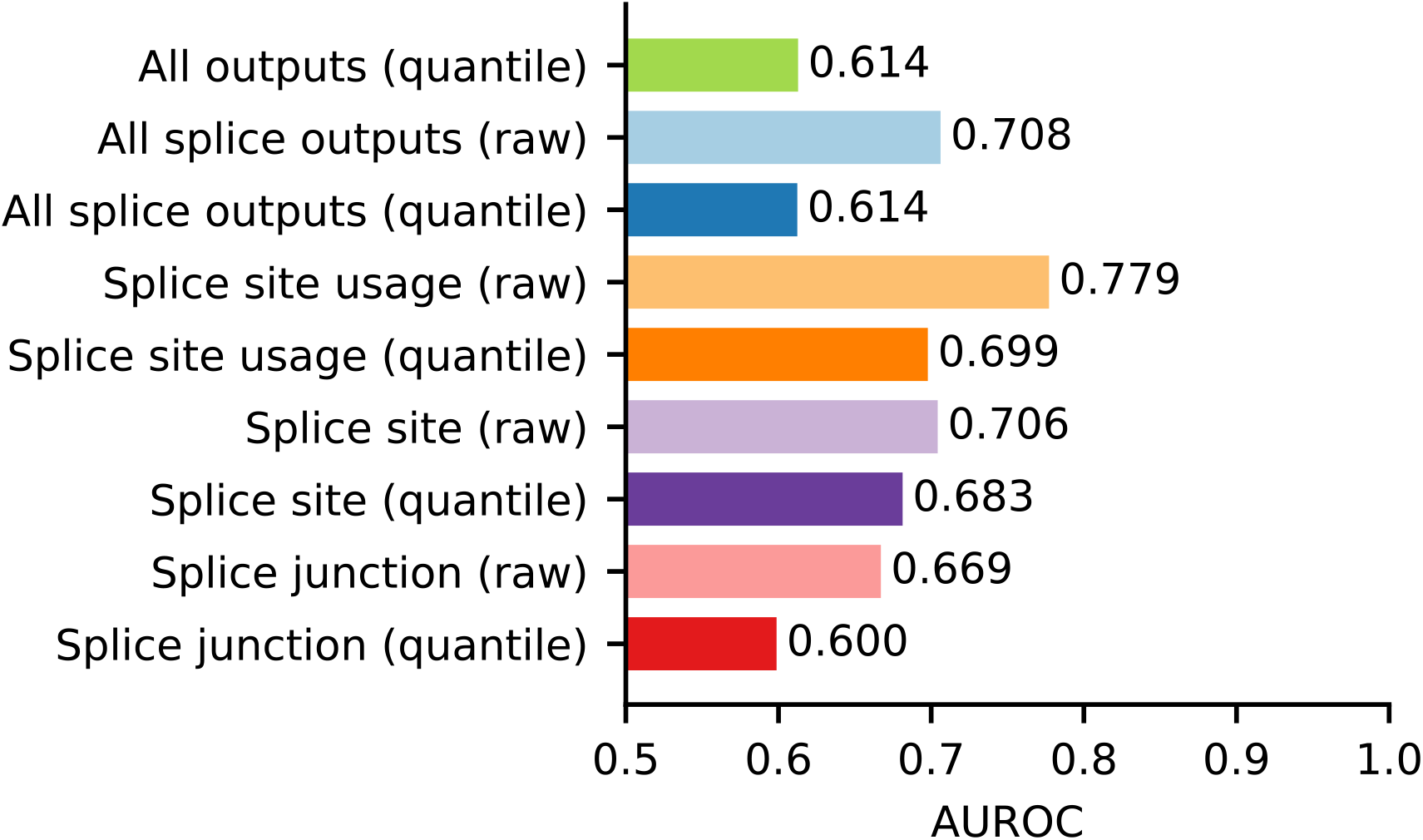
Performance of different AlphaGenome outputs on splice variants. Prediction performance (AUROC) over splice variants (N=11,047, %P=99.32%) using different AlphaGenome outputs. In addition to the composite score integrating all AlphaGenome outputs (as in Figure 1), we also considered specific splicing-related outputs (splice site usage, splice site, and splice junction) and a composite score integrating them, using either raw scores or quantiles (see Methods).

